# Evaluating Tumor Evolution via Genomic Profiling of Individual Tumor Spheroids in a Malignant Ascites from a Patient with Ovarian Cancer Using a Laser-aided Cell Isolation Technique

**DOI:** 10.1101/282277

**Authors:** Sungsik Kim, Soochi Kim, Jinhyun Kim, Boyun Kim, Se Ik Kim, Min A Kim, Sunghoon Kwon, Yong Sang Song

## Abstract

**Background:** Epithelial ovarian cancer (EOC) is a silent but mostly lethal gynecologic malignancy. Most patients present with malignant ascites and peritoneal seeding at diagnosis. In the present study, we used a laser-aided isolation technique to investigate the clonal relationship between the primary tumor and tumor spheroids found in the malignant ascites of an EOC patient. Somatic alteration profiles of ovarian cancer-related genes were determined for eight spatially separated samples from primary ovarian tumor tissues and ten tumor spheroids from the malignant ascites using next-generation sequencing.

**Results:** We observed high levels of intra-tumor heterogeneity (ITH) in copy number alterations (CNAs) and single-nucleotide variants (SNVs) in the primary tumor and the tumor spheroids. As a result, we discovered that tumor cells in the primary tissues and the ascites were genetically different lineages. We categorized the CNAs and SNVs into clonal and subclonal alterations according to their distribution among the samples. Also, we identified focal amplifications and deletions in the analyzed samples. For SNVs, a total of 171 somatic mutations were observed, among which 66 were clonal mutations present in both the primary tumor and the ascites, and 61 and 44 of the SNVs were subclonal mutations present in only the primary tumor or the ascites, respectively.

**Conclusions:** Based on the somatic alteration profiles, we constructed phylogenetic trees and inferred the evolutionary history of tumor cells in the patient. The phylogenetic trees constructed using the CNAs and SNVs showed that two branches of the tumor cells diverged early from an ancestral tumor clone during an early metastasis step in the peritoneal cavity. Our data support the monophyletic spread of tumor spheroids in malignant ascites.

## Background

Epithelial ovarian cancer (EOC) is a silent but mostly lethal gynecologic malignancy. The most common histological EOC subtype is high-grade serous carcinoma, and the current treatment strategy involves a primary debulking surgery followed by chemotherapy to reduce the tumor burden [1, 2]. Recent advances in genomics have revealed the presence of extensive intra-tumor heterogeneity (ITH) in many cancers, including ovarian cancer [3–5]. The presence of extensive clonal diversity increases the capacity of a given tumor to survive upon an expected strike in the microenvironment and thus is thought to be responsible for a reduced response to current chemotherapy and to contribute to chemoresistance development [6–8].

Unlike other solid tumors, the primary route of metastasis in EOC patients is the transcoelomic metastasis route, which is a passive process and involves dissemination of tumor cells from the primary tumor tissue into the peritoneal cavity [9]. Thus, early disseminating clones may exist in the malignant ascites tumor microenvironment (TME) and may form an independent subclonal lineage and contribute to ITH. Both protumorigenic and antitumorigenic factors are known to be enriched in the malignant ascites TME [10]. However, genetic differences between tumor cells in the primary tissue and tumor cells surviving in the ascites TME are not yet fully understood. Multi-region sequencing of both the primary tumor and associated metastases in ovarian cancer has provided insights into spatial heterogeneity and has shown that metastatic tumors maintain the genetic alterations found in the primary tumor and arise with little accumulation of genetic alterations [5]. However, the extent of the genetic heterogeneity within and between the primary tumor and tumor cells found in ascites remains underestimated.

Here, to uncover the genetic heterogeneity of tumor cells in malignant ascites, we introduced a genetic profiling method for individual tumor spheroids which are the common form of tumor cells floating in malignant ascites [11]. Inspired by single-cell analysis, we hypothesized that genetic profiling of individual tumor spheroids might uncover the heterogeneity within and between the primary tumor and tumor cells in ascites. We isolated individual tumor spheroids through a laser-aided isolation technique. Then, we performed low-depth whole-genome sequencing (WGS) and high-depth whole-exome sequencing (WES) for ten tumor spheroids and eight primary tumor samples from a high-grade serous (HGS) EOC patient. We explored somatic copy number alterations (CNAs) and single-nucleotide variants (SNVs) to determine the tumor evolution and ITH between the primary tissues and the tumor spheroids from the malignant ascites. This study reports the feasibility of analyzing tumor cells in malignant ascites to detect early disseminating EOC clones.

## Results

### Preparation and isolation of single tumor spheroids from the ascites of an ovarian cancer patient

A malignant ascites was collected during a primary debulking surgery. The tumor spheroids in the malignant ascites were purified, fixed and prepared on a discharging layer-coated glass slide (Fig. 1A). Single tumor spheroids on the slide were isolated by an infrared (IR) laser pulse as described in our previous publication [12]. Briefly, the discharging layer consisted of indium tin oxide (ITO), which vaporizes when irradiated by an IR laser pulse. The ITO vaporization generates pressure, by which cells in the irradiated area are discharged from the slide. From the prepared sample on the slide, we isolated ten individual tumor spheroids, which were tens of micrometers in diameter and contained hundreds of cells (Fig. 1B, C). Isolating and capturing each tumor spheroid took less than 1 second on average, which means this technique is feasible for analyzing a large number of samples and could be implemented in a routine procedure. The isolated single tumor spheroids were collected in PCR tubes for further reactions.

**Figure 1:**
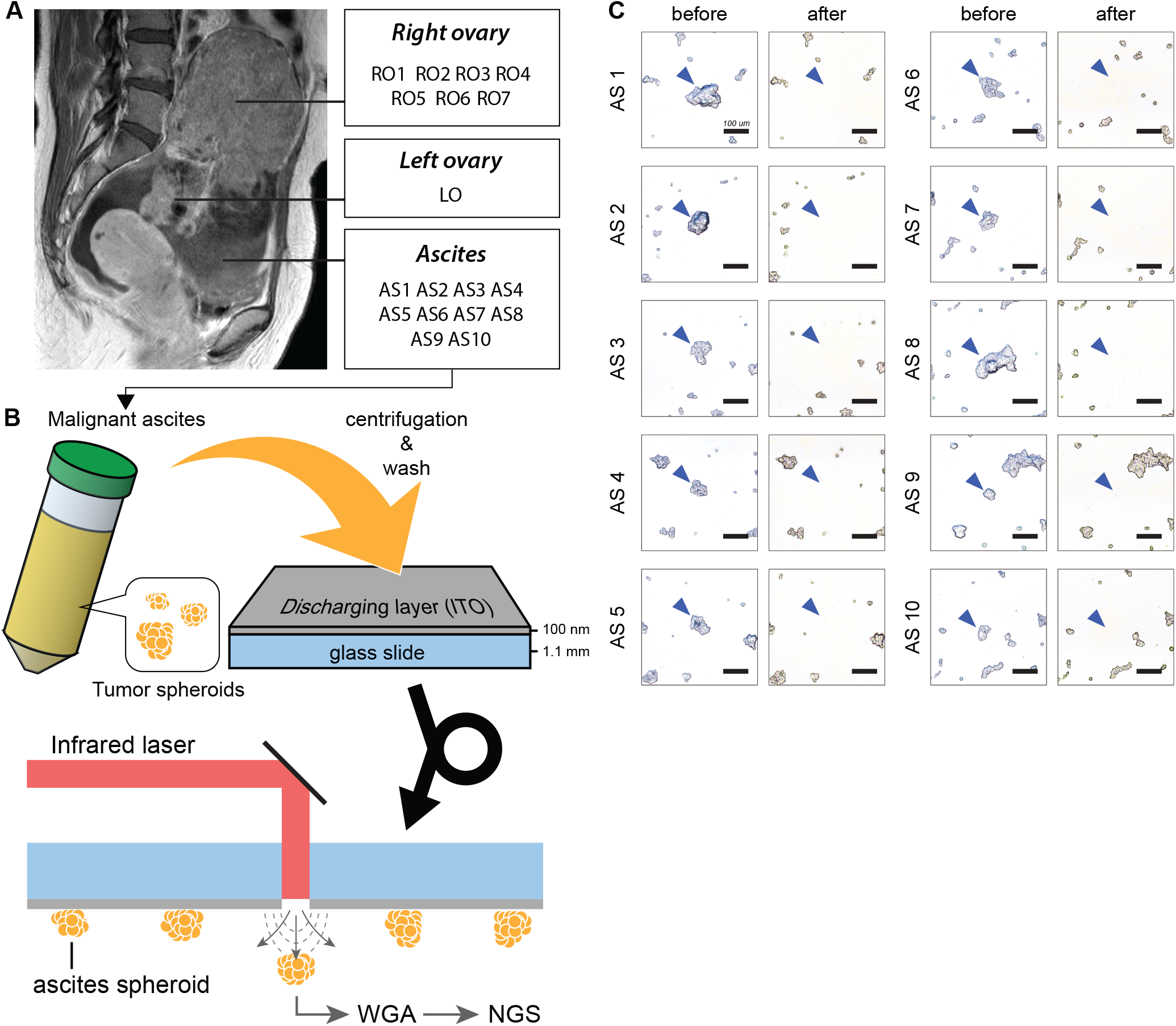
An overview of individual tumor spheroid isolation from malignant ascites. (A) A malignant ascites was collected during a primary debulking surgery. Tumor spheroids in the malignant ascites were purified, fixed and prepared on a discharging layer (Indium Tin Oxide (ITO), 100 nm in thickness)-coated glass slide. (B) The laser isolation technique was used to isolate individual tumor spheroids. This technique utilizes an IR pulsed laser, which vaporizes the discharging layer on the glass slide. Using this technique, ten individual tumor spheroids were isolated from the slide. The isolated cells underwent WGA and sequencing. (C) The images before and after isolation demonstrate that the targeted tumor spheroids in the malignant ascites were specifically isolated without disturbing the neighboring cells. The scale bars represent 100 μm.

### Whole-genome amplification of the isolated individual tumor spheroids

The isolated single tumor spheroids were lysed by proteinase K. Then, the samples underwent multiple displacement amplification (MDA, Fig. 2A). The amplification was monitored via real-time PCR. The results showed that all the isolated samples yielded successful amplification (10/10). Additionally, comparing the amplification plots between the tumor spheroids and controls showed that there was no or a negligible amount of carry-over contamination (Fig. 2B). Every reaction yielded over 2 µg of amplified DNA, which was enough to conduct WGS and WES.

**Figure 2:**
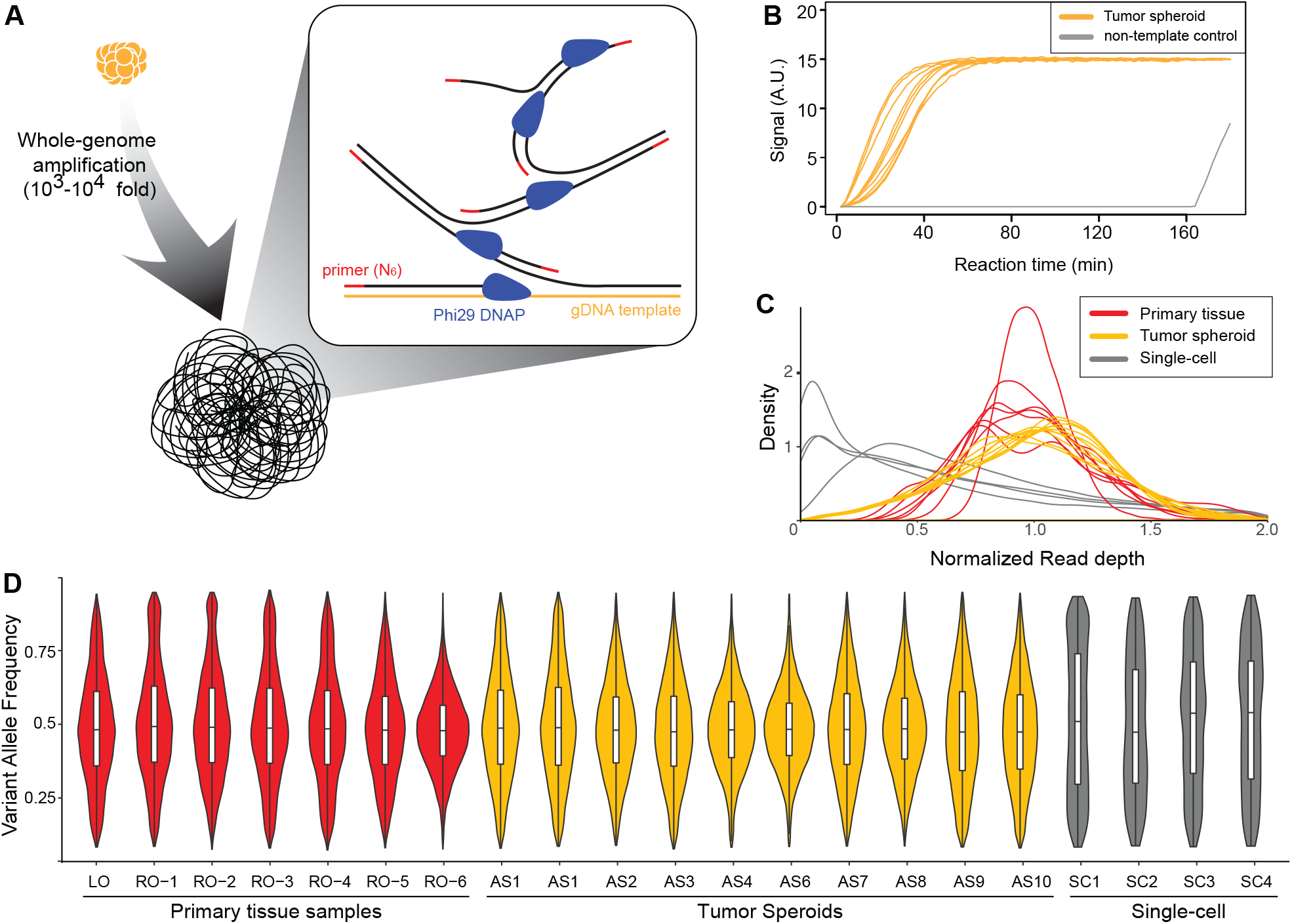
WGA of the isolated tumor spheroids and several quality metrics of the amplified products. (A) MDA was performed to amplify the DNA in each tumor spheroid. MDA amplified tumor spheroid DNA 10^3^-to 10^4^-fold. (B) The amplification process was monitored by observing the fluorescence signal in each reaction. A non-template control was included in the reaction to testify carry-over contamination. The results showed that there was no or a negligible amount of carry-over contamination. (C, D) The distributions of the normalized read depth and VAF reflect the quality of the WGA products. Compared with the distributions of the amplified products from single cells, the distributions of the tumor spheroids were similar to those of the primary tissues. This indicated that the amplified products from the tumor spheroids had a negligible amount of WGA artifacts.

Next, we calculated and plotted the distributions of the normalized read depth (Fig. 2C) and variant allele frequency (VAF, Fig. 2D) based on the sequencing data to evaluate the amplification uniformity of the MDA reaction. In the Fig. 2C and 2D, the distributions of the MDA products from single cells were used for comparison. Normalized read depth indicates the uniformity of the number of sequencing reads throughout the whole-genome. The DNA from bulk tumor samples showed normal-like distributions with small variance, but whole-genome amplified DNA from single cells presented a skewed distribution because of non-uniform amplification. In contrast, the distributions of the tumor spheroids were similar to the distributions of the tumor bulk samples, rather than the whole-genome amplified products from the single cells. This result suggests that the effect of non-uniform amplification during MDA was minimized because hundreds of cells were included in the individual tumor spheroids. Similarly, the VAF distributions of the tumor spheroids were similar to those of the bulk tumor samples but not to the distributions of the single cells. This result supports the presumption that the MDA products of the tumor spheroids present a balanced allele amplification without losing one of the two alleles.

### Low-depth WGS reveals the somatic CNAs and genetic subclones

First, we assessed the somatic CNAs of the primary ovarian cancer tissues and the tumor spheroids from the ascites (Additional file 1: Table S1). We carried out low-depth WGS using the Illumina platform to produce 8.53 ± 0.879 (× 10^6^) sequenced reads for each sample. As a result, we generated CNA profiles based on which we performed a hierarchical clustering analysis (Fig. 3A). The clustering yielded three distinct genetic subgroups. The primary ovarian cancer tissues (RO 1-7 and LO, named “Primary clone” and colored red) were clustered together. In contrast, the tumor spheroids from the ascites were divided into two clusters, one of which showed a primary-like CNA profile (AC 1-3 and 7-8, named “Ascites clone 1” and colored yellow), but the other presented a normal-like profile (AC 4-6 and 9-10, named “Ascites clone 2”, colored green).

**Figure 3:**
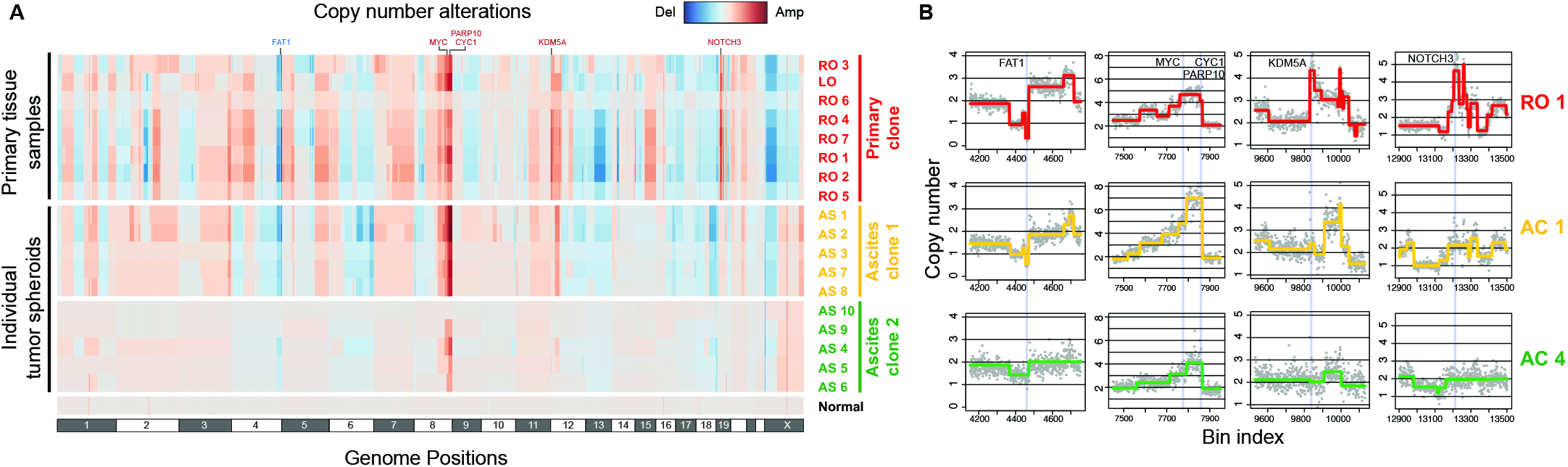
CNA analysis based on the genetic subclones of the tumor cells identified via low-depth WGS. (A) A genome-wide CNA analysis was performed using the low-depth WGS data. Each row represents each sample, and the samples were reordered by the hierarchical clustering method. The clustering analysis generated three major clusters, which were named Primary clone (red), Ascites clone 1 (yellow), and Ascites clone 2 (green). The clear differentiation of the CNA profiles between the Primary clone and Ascites clones implied that the tumor spheroids in the Ascites clones were not derived from the tumor cells in the Primary clone but from another independent tumor lineage. (B) Representation of the CNA profiles in detail at several regions for RO1, AC1, and AC4. The three samples exhibited both shared and exclusive CNAs. For example, deletion of FAT1 (1^st^ column) and amplification of MYC, CYC1, and PARP10 (2^nd^ column) were shared in every sample. However, the amplification of KDM5A (3^rd^ column) and NOTCH3 (4^th^ column) was exclusive to the Primary clone. This might indicate that the FAT1, MYC, CYC1, or PARP10 alterations conferred a growth advantage to the common ancestor of the Primary clone and Ascites clones. In contrast, the KDM5A or NOTCH3 amplifications might cause branching from the common ancestor and proliferation of the Primary clone

Interestingly, the CNA profiles showed that deletion of FAT1 and amplification of MYC, PARP10, and CYC1 were shared by most of the samples (Fig. 3B). These genes are reported to be recurrently deleted (FAT1) or amplified (MYC, PARP10, and CYC1) in pan-cancer data [13]. These facts suggest that the shared CNAs might be the driving alterations at the first stage of cancer initiation. However, the primary clone had exclusive focal amplifications of KDM5A and NOTCH3 (Fig. 3B), which are known as recurrently amplified genes in ovarian cancer [13, 14]. These focal amplifications of KDM5A and NOTCH3 might allow the primary clone to overwhelm the other subclones and finally dominate the left and right ovaries. However, we did not find a critical focal amplification or a deep deletion exclusive to Ascites clone 1. This implied that other types of alterations might drive Ascites clone 1 to survive or propagate in the peritoneal fluid.

### WES reveals somatic SNVs and genetic subclones

To identify the somatic SNVs, the samples underwent WES. For each sample, the sequencing run generated 134 ± 21.4 depth of data, covering the whole exome of the human genome. As a result, 171 somatic SNVs were identified by variant calling from all the samples (Additional file 2: Table S2). The results shown in Fig. 4A and 4B revealed that 38.6% of the SNVs were common to the primary tumor and tumor spheroids from the ascites, and 35.7% of the SNVs exclusively belonged to primary-only and 25.7% to ascites-only mutations. The exclusive mutations in the Ascites clone suggest that this clone evolved by accumulating mutations independent from the Primary clone. Interestingly, the Ascites clone had a nonsynonymous mutation in the KRAS gene at 12p12.1. A single nucleotide substitution (C>T) results in an activating KRAS mutation that is a well-known oncogenic mutation associated with the anchorage-independent growth of tumor cells through the acquisition of anoikis resistance in various malignancies [15, 16]. Therefore, the mutation in KRAS in the Ascites clone might provide an additional fitness gain for anchorage-independent survival in the ascites TME. However, both the Primary and Ascites clones shared somatic SNVs in TP53 and ARID1A, which are well-known driver mutations in ovarian cancer [17, 18]. At the initial stage of tumorigenesis, these mutated genes might be tumor-initiating SNVs in conjunction with the CNAs of FAT1, MYC, PARP10, and CYC1. In addition to these somatic variants, the patient had germline variants in BRCA1 (NM_007294.3:c.1511dupG) and TP53 (NM_001126118:c.C98G), which are well-known susceptibility genes of ovarian cancer and are likely to predispose individuals to ovarian cancer and promote carcinogenesis (Additional file 3: Table S3) [19, 20].

**Figure 4:**
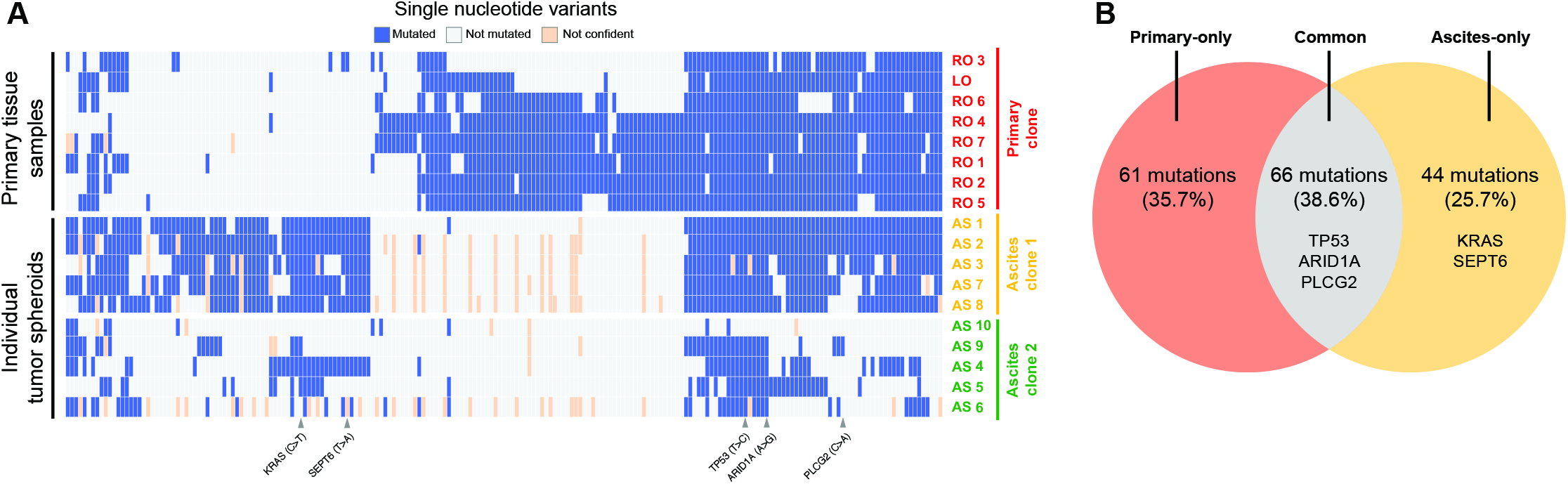
SNV analysis based on the WES data. The WES data from the primary tissue samples and tumor spheroids were used to analyze the SNVs. The results showed that a significant portion of the SNVs was shared in the Primary clone and Ascites clone 1. At the same time, the Primary clone and Ascites clone 1 had unique mutations. This result suggests that the two clones might have branched from a common ancestor. Ascites clone 2 was excluded from the analysis because the tumor spheroids in Ascites clone 2 were presumed to contain a large number of normal cells in each tumor spheroid.

### Cellular composition of the tumor spheroids

Regarding the CNAs, Ascites clone 2 had no alteration except for amplification of the 8q24 region. Concerning the SNVs, Ascites clone 2 had fewer mutations than the other clusters. Based on these facts, we examined the possibility that normal cells exist in a tumor spheroid. We assumed that the VAF distribution of Ascites clones 1 and 2 would be similar if the two subclones had a similar proportion of normal cells. However, the VAF of Ascites clone 2 would be low if a single tumor spheroid from the clone included a high proportion of normal cells. We tested this idea by plotting the VAF distribution of each sample (Fig. 5). The results showed that most of the VAF distributions from the Primary clone and Ascites clone 1 were located at a higher range than those from Ascites clone 2. Therefore, we concluded that the small number of CNAs and SNVs in Ascites clone 2 was not due to their true characteristics but because the proportion of tumor cells in the tumor spheroid was small. Consequently, we excluded Ascites clone 2 from the following phylogenetic analysis.

**Figure 5:**
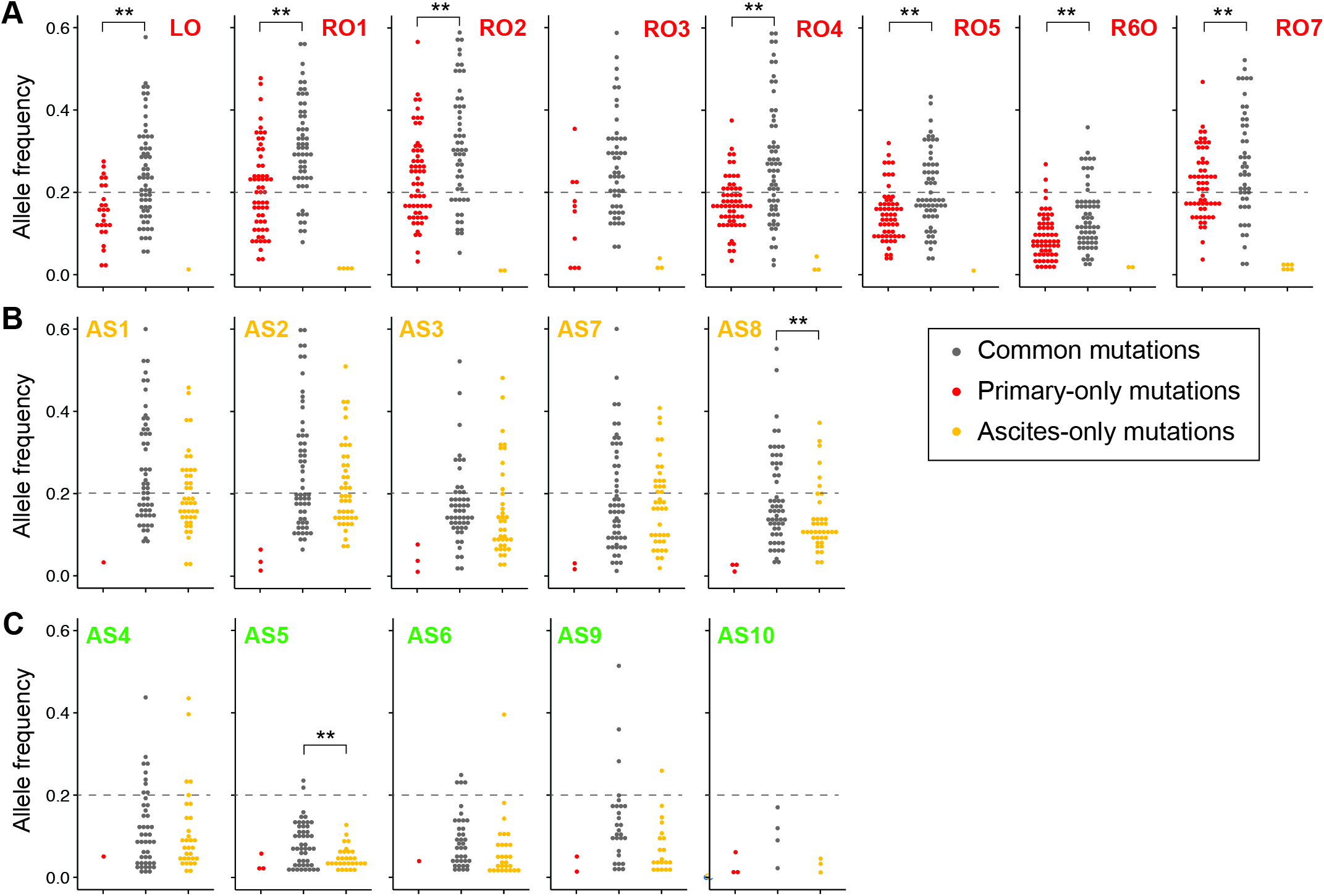
Analysis of the allele frequency to infer the cellular composition of each sample. The VAF distribution was plotted for each sample from the (A) Primary site and (B, C) Ascites. The mutations were categorized into common, primary-only or ascites-only mutations. Common mutations were somatic SNVs, which were detected in both the Primary clone and Ascites clones, and primary-only and ascites-only mutations, which were shared somatic SNVs detected only in the Primary clone and Ascites clones, respectively. The results showed that most of the VAF distributions from Ascites clone 2 were located at a much lower range than those from the Primary clone and Ascites clone 1. This suggests that the tumor spheroids in Ascites clone 2 had a large proportion of normal cells in each tumor spheroid.

In addition to the presence of normal cells in the samples, we examined the possibility of the presence of heterogeneous tumor cells in the samples. By comparing the allele frequency distributions of the common and primary-only mutations for each sample, we found that the allele frequencies of the common mutations were higher than those of the primary-only mutations for the primary tissues (7 of 8 samples, p < 0.01). This result implies that each of the primary tissues (except RO3) had two or more subclones sharing common mutations but not subclonal mutations. In contrast, the allele frequencies of the common mutations were similar to those of the ascites-only mutations for the tumor spheroids (8 of 10 samples). This result can be interpreted to indicate that, compared with the primary tissue samples, each tumor spheroid was comprised of genetically homogeneous tumor cells.

### Constructing phylogenetic trees based on the somatic CNAs and SNVs

The phylogenetic trees were constructed from the CNA and SNV data. We achieved a CNA-based phylogeny analysis by identifying the common chromosomal breakpoints, calculating a trinary event matrix, and constructing a maximum parsimony tree [28]. The phylogenetic tree showed that an ancestral cancer clone accumulated CNAs and divided into two clones, which gained additional exclusive CNAs (Fig. 6A). Notably, these two genetic clones were composed of tumor spheroids from ascites and tumor tissues. Potentially, physically separated and biologically distinct TMEs might drive cancer cells into different alteration statuses.

**Figure 6:**
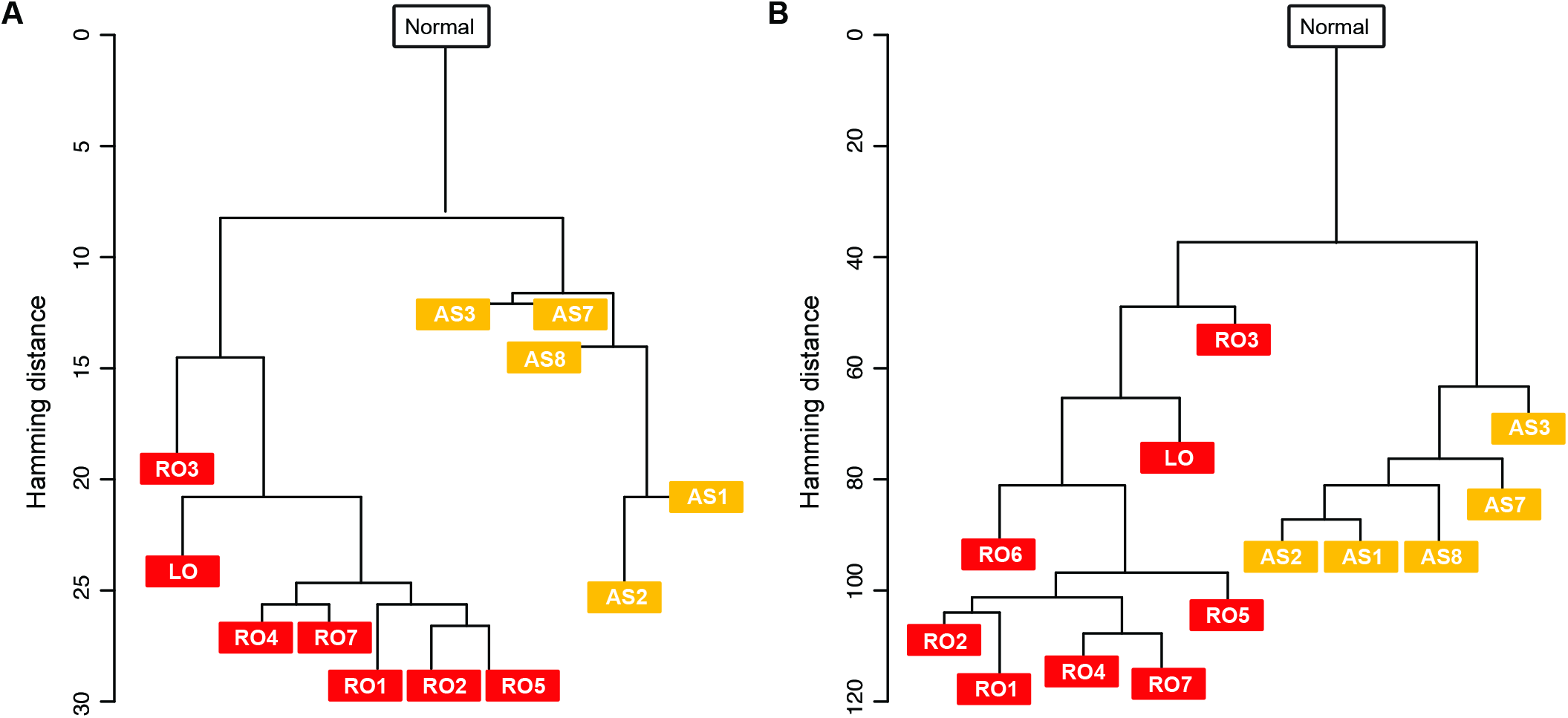
Constructing phylogenetic trees. Phylogenetic trees were constructed using both the (A) CNA profiles and (B) SNV profiles. The two trees presented similar topologies and indicated that the Primary clone and Ascites clone 1 were derived from one ancestral clone at the early stage of cancer development. In addition, the phylogenic trees indicated that the analyzed tumor spheroids were not derived from the primary tumor cells that were present at the time of sampling.

Maximum parsimony tree generation using the CNA data has a couple of limitations. First, this approach needs to set thresholds to define the amplified, neutral, and deleted status. The resultant tree is significantly affected by thresholds, and there is no golden rule to set the thresholds. Second, the proportion of normal cells in a sample has a substantial impact on a tree because the CNA status might be incorrectly assigned according to the normal cell portion. For example, the VAFs of RO6 (Fig. 5) show that the sample had a large number of normal cells. In this case, the copy number value of RO6 was close to the normal value (Fig. 3A), although the overall pattern was not similar to that of the normal sample. Thus, the thresholding led RO6 to be the same as the normal sample. For this reason, we excluded RO6 when constructing the maximum parsimony tree based on the CNA data.

Next, we constructed a phylogenetic tree from the SNV data. This approach does not use manual thresholding, and a phylogenetic tree is less affected by a normal cell portion. Therefore, we expected that, compared with the CNA-based approach, this approach would provide a more accurate result. The results showed that the cancer cells accumulated mutations as a single clone and divided into two independent clones (Fig. 6B). Moreover, with the full advantage of the SNV information, the phylogenetic tree presented the sequential creation of RO3, LO, and the rest of the Primary clones. Overall, the phylogenetic tree based on the SNV data rather than the CNA data presented a more stable and biologically explainable result.

### Inferring the evolutionary trajectory of the primary ovarian cancer and the single tumor spheroids in the ascites

This patient harbored a bilateral ovarian tumor at the time of the primary debulking surgery. It is important to note whether these bilateral tumors arise independently or are the result of metastasis. The clonal evolution of the tumorigenesis theory provides two mechanisms of bilateral ovarian tumor development. If bilateral ovarian tumors arise from independent ancestral clones, they would have distinct genomic profiles without sharing somatic alterations. In contrast, bilateral tumors would have an identical set of somatic variants if they resulted from metastasis [21]. The somatic CNAs and SNVs of the left and right primary ovarian tumor in this study displayed comparable genomic profiles, strongly indicating a monoclonal origin of the bilateral tumor in this patient. This was further confirmed by calculating the clonality index (CI) based on previous reports [21, 22] revealing that the bilateral ovarian tumors were clonally related (CI1 = 1.0).

Finally, the history of the ovarian cancer development and progression was established based on the genomic profiles to understand the tumor evolution and its direction in this patient. As noted earlier, ovarian cancer metastasis occurs through a passive process, which initially involves physical shedding of tumor cells from the primary tumor into the peritoneal cavity, and the accumulation of ascites facilitates distant seeding of tumor cells along the peritoneal wall. Given a fixed chance of evolution, two scenarios are possible, either a monoclonal or polyclonal seeding process. If only certain clones from the primary tumor are fit to survive in the ascites TME, distinct clones, which may have diverged early, may be selected and progress over time in the primary and ascites TMEs, showing a tendency toward independent tumor evolution driven by different TMEs. In contrast, if tumor evolution is entirely driven by clonal dominance and the physical shedding of tumor cells from the primary tumor occurs by chance, then dominant clones expand in size and others may remain unchanged or become extinct over time at the primary tumor site. As the tumor grows, multiple clones may shed from the primary tumor into ascites. The ascites TME then acts as a reservoir of clonal lineage, and tumor cells in the ascites would represent the entire mutational landscape of a given tumor. For our case, we observed significant genetic differences in the CNAs and SNVs among the primary tissue samples and tumor spheroids. The dominant clones found in the right ovary were absent in the ascites TME, and we found 44 tumor spheroid–specific somatic SNVs (Additional file 2: Table S2). Furthermore, the comparable allele frequencies between the common mutations and tumor spheroid–specific mutations suggest that the tumor spheroids in the ascites TME are comprised of genetically homogeneous tumor cells compared with the primary tissues. Therefore, we conclude that the tumor spheroids were from a single subclonal lineage, supporting a mono-and early-seeding origin of the tumor spheroids in this patient. Based on these perspectives, we drew a potential evolutionary trajectory of the tumor from the patient (Fig. 7A). The tumor was initiated at the right ovary to generate the ancestral clone. With further accumulation of mutations, the ancestral clone evolved into two subclones, the first of which was found in the right ovary and metastasized to the left ovary. The second subclone shed into the ascites TME and became extinct or dominated by the first subclone in the right ovary (Additional file 4: Table S4). Eventually, the Ascites subclone moved to the peritoneal cavity. In addition, the summary of genome-wide somatic CNAs and SNVs indicated that the tumor cells in the primary tissue and the ascites possessed exclusive alterations as well as common ones (Fig. 7B). This result shows that the tumor cells in the primary tissue and the ascites were two subclonal lineages, which branched from one ancestral lineage.

**Figure 7:**
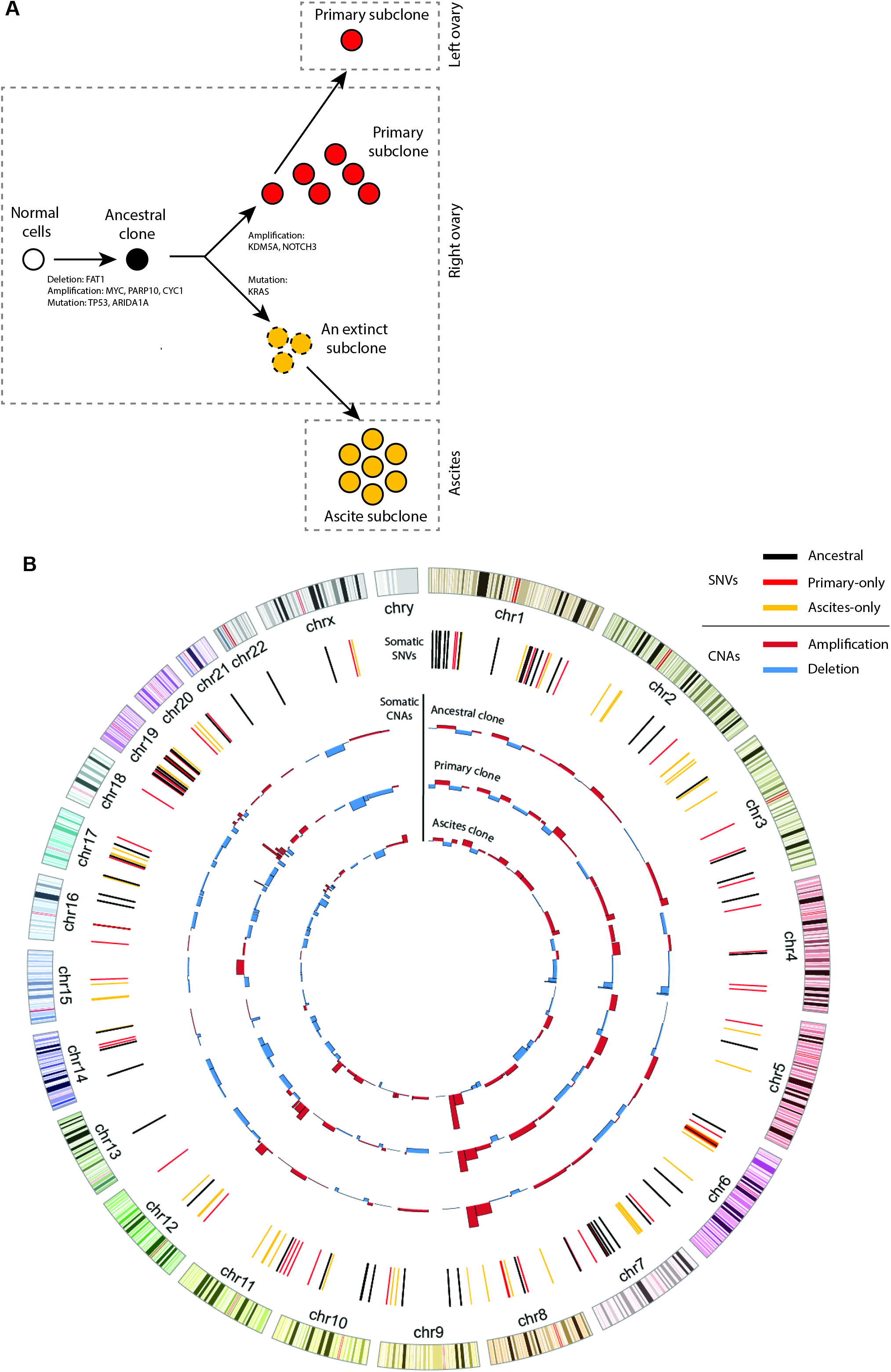
The inferred evolutionary history of the tumor and the Circos plot of the major subclones. (A) Based on the sequencing data from the primary tissue samples and the tumor spheroids from the ascites, the evolutionary trajectory was inferred. The tumor was initiated at the right ovary to generate the ancestral clone. With the further accumulation of mutations, the ancestral clone evolved into two subclones, the first of which was found in the right ovary and metastasized to the left ovary. The second subclone shed into the ascites TME and became extinct or dominated by the first subclone in the right ovary. Eventually, the Ascites subclone moved to the peritoneal cavity. (B) The Circos plot presents the genome-wide alterations in the ancestral, Primary, and Ascites subclones. For the SNVs, the black, red, and yellow bars represent the ancestral, primary-only, and ascites-only mutations, respectively. The tumor cells acquired the ancestral mutations before dividing into the Primary and Ascites clones. After division, the Primary and Ascites clones acquired lineage-specific SNVs. For the CNAs, the red and blue bars represent amplification and deletion, respectively.

## Discussion

In this study, we attempted to determine the presence of genetic heterogeneity within and between a primary tumor and the associated tumor spheroids in the ascites by performing multi-region sequencing of the primary tumor and genetic profiling of the individual tumor spheroids using the laser-aided cell isolation technique. We performed both WGS and WES of the primary tumor and tumor spheroid samples. First, we discovered high ITH levels in eight primary tissues and ten tumor spheroids. We also discovered that the CNA profiles in the primary and associated tumor spheroids were separated into two distinct genetic clusters, suggesting that the TME may be operative during tumor evolution. Second, we identified somatic SNVs using WES. We discovered a total of 171 somatic SNVs from all the samples, and 66 (38.6 %) of these SNVs were ubiquitous mutations that were common to the primary tumor and tumor spheroids. The rest were either primary-only (61 SNVs, 35.7 %) or ascites-only (44 SNVs, 25.7 %) mutations, highlighting the notion that the tumor spheroids might have diverged early and accumulated additional mutations independently from the Primary clone. Supporting this idea, both phylogenic analyses, using the CNAs and SNVs, showed that the tumor spheroids might have diverged early from an ancestral tumor clone, evolved further with distinctive genomic profiles, and formed an independent subclonal lineage, thereby contributing to the ITH.

We also assessed the normal cell contamination in both the primary tumor and tumor spheroids using the VAF distribution in each sample. Indeed, both the Primary clone and Ascites clone 1 showed higher VAF distributions than Ascites clone 2, suggesting that the normal-like CNA and SNV profiles in Ascites clone 2 were due to a high proportion of normal cells. These findings are consistent with previous data from ovarian cancer patient-derived tumor spheroids and mouse models that suggested the presence of tumor-associated macrophages in the center of tumor spheroids [23].

Although we only studied a single high-grade EOC patient, our data support previous studies demonstrating early divergence of the ascites sample from the primary tumor [24]. Further studies are needed to compare similarities and differences between the ascites spheroids and distant metastasis samples. Our data suggest that the mutation set of ascites spheroids does not represent the entire mutational landscape of a given EOC patient. This disagrees with recent findings by Choi et al. [25] showing that ascites tumor cells represent the entire mutational landscape of a given tumor, and no additional genetic aberrations were detected. In contrast, our data showed the presence of genetic heterogeneity within and between the primary tumor and the associated ascites spheroids. Moreover, the primary and associated ascites spheroids diverged early in tumor development, and not all the Primary clones disseminated into the ascites TME. However, our study is limited to a single ascites TME and provided no insight into distant metastatic sites.

Additionally, our data demonstrated that, compared with the primary tissue samples, each tumor spheroid was comprised of genetically homogeneous tumor cells (Fig. 5). This can be interpreted in two ways. First, the tumor cells in an ascites may have low ITH. In this case, the spheroids of the tumor cells would be genetically homogeneous. Second, the tumor cells with a similar genetic profile may form individual tumor spheroids. In this case, the tumor cells in each tumor spheroid might have the same genetic profile, but two different tumor spheroids might be genetically different. For this case, isolating and analyzing the individual tumor spheroids from ascites might be widely utilized to discover the ITH of ovarian cancer.

Our data can partly be explained by the theory of Darwinian selection. For simplicity, tumor evolution is described as a series of expansions of clones, where each expansion series is driven by additional mutation acquisition, and clone fitness is tested by Darwinian selection. This selective sweep is context-dependent, and thus, genetic variants that are beneficial at a certain point may become extinct throughout the period of tumor progression. As a consequence, these clones may be absent in a fully grown tumor [26]. The selective pressures are further influenced by the dynamics of the TME, thereby increasing the complexity of tumor evolution [27]. The presence of extensive ITH in tumor spheroids and the early divergence of these subclones from the primary tumor suggests that we are currently underestimating the tumor genomic landscape.

In addition to the importance of genetic differences between tumor cells in primary tissue and those in ascites, knowledge regarding the genetic heterogeneity within the tumor cells in ascites would be valuable. Although not thoroughly studied, the genetic diversity of tumor cells in an ascites may have a large impact on tumor relapse and metastasis, given that transcoelomic spread is the primary route of metastasis in ovarian cancer. However, there has been no attempt to discover the genetic heterogeneity of individual tumor spheroids. In this study, we evaluated 10 individual tumor spheroids, five of which contained sufficient tumor cells for the analysis. Although we observed genetic heterogeneity of the individual ascites spheroids, a follow-up study should analyze at least a few tens of individual tumor spheroids per patient to find a clear signature of the genetic heterogeneity in an ascites.

## Conclusion

In this study, we performed genome-wide sequence analysis of the primary tumor and the associated tumor spheroids in the malignant ascites of an EOC patient. We analyzed genetic heterogeneity in the primary tumor and tumor spheroids through multi-region sequencing and the laser-aided cell isolation technique [12]. From the sequencing data, we discovered clonal or subclonal somatic CNAs and SNVs, based on which we constructed phylogenetic trees and inferred the evolutionary history of tumor cells in the patient. As a result, we found that the tumor cells in the malignant ascites were an independent lineage from the primary tumor. The phylogenetic analysis showed that the lineage branched before the evolution of the cancer cells at the primary tissues, which suggests that analyzing malignant ascites might be used to detect ovarian cancer or metastasis in the early stage. In summary, the genetic plasticity and similarity between a primary tumor and associated tumor spheroids are still not clear, and yet, the nature of the similarity may have profound implications for both tumor progression and therapeutic outcomes in ovarian cancer. Therefore, future prospective studies profiling the genomic information of primary ovarian tumors, distant metastatic tumors, and tumor spheroids to determine the direction of tumor evolution and metastasis of ovarian cancer are warranted.

## Methods

### Patient information and sample preparation

A 42 yr old female patient diagnosed with primary high-grade serous ovarian cancer (Grade 3, stage IIIC) presented with malignant ascites and peritoneal seeding. Both primary tissues and malignant ascites were collected during primary debulking surgery. Fresh primary tissues and tumor cell clusters were mounted onto ITO-coated glass slides. Six samples were taken randomly from the solid portions of right ovary and only one from left ovary. Blood was collected to serve as the normal control. Ten tumor cell clusters were collected from the malignant ascites and fixed in 10% (v/v) formaldehyde. This study was approved by the Institutional Review Board (IRB) at Seoul National University Hospital (Registration number: 1305-546-487) and performed in compliance with the Helsinki Declaration. We obtained informed consent from the patient prior to primary debulking surgery to be used in research.

### Laser-aided isolation of tumor spheroids and their whole-genome amplification

Previously, we developed and published a laser-aided cell isolation technique [12], and designed two different pieces of software written in Python scripts and available at Github (https://github.com/BiNEL-SNU/PHLI-seq). Isolation of tumor spheroids was performed as described in the prior publication. In brief, an infrared laser was applied to the target area, vaporizing Indium Tin Oxide (ITO) layer and discharging the targeted tumor spheroid on the region. We used glass slides with a 100-nm-thick ITO layer.

The 8-strip PCR tube caps for the retrieval of tumor spheroids were pre-exposed under O_2_ plasma for 2 minutes. The tumor spheroids were lysed using proteinase K (cat no. P4850-1ML, Sigma Aldrich) according to the manufacturer’s directions after the PCR tubes were centrifuged. For whole-genome amplification, we used GE’s Illustra Genomiphi V2 DNA amplification kit (cat no. 25-6600-30). We added 0.2 µl of SYBR green I (Life Technologies) into the reaction solution for real-time monitoring of the amplification (Fig. 2B). All amplified products were purified using Beckman Coulter’s Agencourt AMPure XP kit (cat no. A63880) immediately following the amplification. To prevent carry-over contamination, the pipette tip, PCR tube, and cap for the reaction were stored in a clean bench equipped with UV light and treated with O_2_ plasma for 2 minutes before use. Additionally, we monitored the real-time amplification of non-template controls to ensure that no contaminants were transferred.

### Sequencing library preparation, whole-genome, and whole-exome sequencing

The whole-genome amplified products or genomic DNA were fragmented using an EpiSonic Multi-Functional Bioprocessor 1100 (Epigentek) to generate DNA fragments with 250-bp on average. The fragmented products underwent Illumina library preparation using Celemics NGS Library Preparation Kit (LI1096, Celemics, Seoul, Korea) for the whole-genome sequencing library preparation, and SureSelectXT (Agilent, CA, US) for whole-exome sequencing. DNA purification was performed by TOPQXSEP MagBead (XB6050, Celemics, Seoul, Korea), and DNA libraries were amplified using the KAPA Library Amplification Kit (KAPA Biosystems, KK2602). Finally, the products were quantified by TapeStation 2200 (Agilent, CA, US). We used HiSeq 2500 150 PE (Illumina) to generate 1 Gb/sample for whole-genome sequencing and 5 Gb/sample for whole-exome sequencing, respectively.

### Detecting copy number alterations

We used low-depth whole-genome sequencing data and the variable-size binning method [29] to estimate the CNAs of the samples. Briefly, the whole genome was divided into 15,000 variable-sized bins (median genomic length of bin = 184 kbp), in which each bin had an equal expected number of uniquely mapped reads. Then, each sequence read was assigned to each bin followed by Lowess GC normalization to obtain the read depth of each bin. The copy number was estimated by normalizing the read depth of each bin by the median read depth of the reference DNA.

### Detecting Single Nucleotide Variants

GATK (v3.5-0) IndelRealigner and BaseRecalibrator were used to locally realign reads around the Indel and recalibrate the base quality score of BAM files [30]. Then, GATK UnifedGenotyper, Varscan, and MuTect were used and combined the results to avoid false-positive variant calls [31]. First, GATK UnifiedGenotyper was used with default parameters followed by GATK VariantRecalibrator to obtain filtered variants [30]. Data of primary tissue samples and ascites tumor spheroid samples were processed together to produce a single vcf file. dbSNP build 137, HapMap 3.3, Omni 2.5, and 1000G phase1 were used as the training data for variant recalibration. Also, annotation data including QD, MQ, FS, ReadPosRankSum, and MQRankSum were used for the training. Variants detected in the paired blood sample of the cancer patient were removed to produce the final list of GATK called variants. Varscan2 [32] (ver 2.3.7) and Mutect [33] (ver 1.1.4) were used with default parameters to produce the lists of Varscan and MuTect called variants, respectively. Here, paired blood read data was also used to remove germline variants.

Among the variants from the three callers, variants called by at least two callers were collected to obtain intra-sample double-called sites. We could reduce false-positive variant caused by NGS errors by considering only double-called variants for subsequent analysis [31]. Among the intra-sample double called sites, variants found in at least two samples were collected to remove WGA (whole genome amplification) errors, and the genomic loci with the resultant variants were considered confident sites. Finally, a variant in the confident sites was considered to be true if one of the three variant callers detected the variant at the locus and the allele count of the variant was significantly larger than that of the other non-reference bases (Fisher’s exact test, p < 10^-3^). The overall process is visually described in Additional file5: Figure S1.

## Abbreviations

EOC: Epithelial ovarian cancer
ITH: Intra-tumor heterogeneity
CNA: Copy number alteration
SNV: Single-nucleotide variant
TME: Tumor microenvironment
WGS: Whole-Genome Sequencing
WES: Whole-exome sequencing
HGS: High-grade serous
ITO: Indium tin oxide
MDA: Multiple displacement amplification
VAF: Variant allele frequency
CI: Clonality index

## Declarations

### Ethics approval and consent to participate

This study was approved by the Institutional Review Board (IRB) at Seoul National University Hospital (Registration number: 1305-546-487) and performed in compliance with the Helsinki Declaration. We obtained informed consent from the patient prior to primary debulking surgery to be used in research.

### Consent for publication

Not applicable

### Availability of data and materials

The datasets used and/or analyzed during the current study are available from the corresponding author on reasonable request

### Competing interests

The authors declare that they have no competing interests

## Acknowledgements

Not applicable

## References

1. Reid BM, Permuth JB, Sellers TA. Epidemiology of ovarian cancer: a review. Cancer Biol Med. 2017;14:9–32.

2. Jessmon P, Boulanger T, Zhou W, Patwardhan P. Epidemiology and treatment patterns of epithelial ovarian cancer. Expert Rev Anticancer Ther. 2017;17:427–37.

3. Ding L, Ley TJ, Larson DE, Miller CA, Koboldt DC, Welch JS, et al. Clonal evolution in relapsed acute myeloid leukaemia revealed by whole-genome sequencing. Nature. 2012;481:506–10.

4. Gerlinger M, Rowan AJ, Horswell S, Math M, Larkin J, Endesfelder D, et al. Intratumor heterogeneity and branched evolution revealed by multiregion sequencing. N Engl J Med. 2012;366:883–92.

5. Lee JY, Yoon JK, Kim B, Kim S, Kim MA, Lim H, et al. Tumor evolution and intratumor heterogeneity of an epithelial ovarian cancer investigated using next-generation sequencing. BMC Cancer. 2015;15:85.

6. Swanton C. Intratumor heterogeneity: evolution through space and time. Cancer Res. 2012;72:4875–82.

7. Bilen MA, Hess KR, Campbell MT, Wang J, Broaddus RR, Karam JA, et al. Intratumoral heterogeneity and chemoresistance in nonseminomatous germ cell tumor of the testis. Oncotarget. 2016;7:86280–9.

8. Greaves M. Evolutionary determinants of cancer. Cancer Discov. 2015;5:806–20.

9. Tan DS, Agarwal R, Kaye SB. Mechanisms of transcoelomic metastasis in ovarian cancer. Lancet Oncol. 2006;7:925–34.

10. Kim S, Kim B, Song YS. Ascites modulates cancer cell behavior, contributing to tumor heterogeneity in ovarian cancer. Cancer Sci. 2016;107:1173–8.

11. Smolle E, Taucher V, Haybaeck J. Malignant ascites in ovarian cancer and the role of targeted therapeutics. Anticancer Res. 2014;34:1553–61.

12. Kim S, Lee AC, Lee H, Kim J, Jung Y, Ryu HS, et al. Constructing and visualizing cancer genomic maps in 3D spatial context by Phenotype-based High-throughput Laser-aided Isolation and Sequencing (PHLI-seq). bioRxiv 2018; doi: 10.1101/278010

13. Zack TI, Schumacher SE, Carter SL, Cherniack AD, Saksena G, Tabak B, et al. Pan-cancer patterns of somatic copy number alteration. Nat Genet. 2013;45:1134–40.

14. Park JT, Li M, Nakayama K, Mao TL, Davidson B, Zhang Z, et al. Notch3 gene amplification in ovarian cancer. Cancer Res. 2006;66:6312–8.

15. Derouet M, Wu X, May L, Hoon Yoo B, Sasazuki T, Shirasawa S, et al. Acquisition of anoikis resistance promotes the emergence of oncogenic K-ras mutations in colorectal cancer cells and stimulates their tumorigenicity in vivo. Neoplasia. 2007;9:536–45.

16. Rytomaa M, Martins LM, Downward J. Involvement of FADD and caspase-8 signalling in detachment-induced apoptosis. Curr Biol. 1999;9:1043–6.

17. Ahmed AA, Etemadmoghadam D, Temple J, Lynch AG, Riad M, Sharma R, et al. Driver mutations in TP53 are ubiquitous in high grade serous carcinoma of the ovary. J Pathol. 2010;221:49–56.

18. Wiegand KC, Shah SP, Al-Agha OM, Zhao Y, Tse K, Zeng T, et al. ARID1A mutations in endometriosis-associated ovarian carcinomas. N Engl J Med. 2010;363:1532–43.

19. Welcsh PL, King MC. BRCA1 and BRCA2 and the genetics of breast and ovarian cancer. Hum Mol Genet. 2001;10:705–13.

20. Xi Y, Liu C, Xin X. Association between a single nucleotide polymorphism in the TP53 region and risk of ovarian cancer. Cell Biochem Biophys. 2014;70:1907–12.

21. Yin X, Jing Y, Cai MC, Ma P, Zhang Y, Xu C, et al. Clonality, heterogeneity and evolution of synchronous bilateral ovarian cancer. Cancer Res. 2017;77:6551–61.

22. Schultheis AM, Ng CK, De Filippo MR, Piscuoglio S, Macedo GS, Gatius S, et al. Massively parallel sequencing-based clonality analysis of synchronous endometrioidendometrial and ovarian carcinomas. J Natl Cancer Inst. 2016;108:djv427.

23. Yin M, Li X, Tan S, Zhou HJ, Ji W, Bellone S, et al. Tumor-associated macrophages drive spheroid formation during early transcoelomic metastasis of ovarian cancer. J Clin Invest. 2016;126:4157–73.

24. Schwarz RF, Ng CK, Cooke SL, Newman S, Temple J, Piskorz AM, et al. Spatial and temporal heterogeneity in high-grade serous ovarian cancer: a phylogenetic analysis. PLoS Med. 2015;12:e1001789.

25. Choi YJ, Rhee JK, Hur SY, Kim MS, Lee SH, Chung YJ, et al. Intraindividual genomic heterogeneity of high-grade serous carcinoma of the ovary and clinical utility of ascitic cancer cells for mutation profiling. J Pathol. 2017;241:57–66.

26. Marusyk A, Polyak K. Tumor heterogeneity: causes and consequences. Biochim Biophys Acta. 2010;1805:105–17.

27. Polyak K, Haviv I, Campbell IG. Co-evolution of tumor cells and their microenvironment. Trends Genet. 2009;25:30–8.

28. Gao R, Davis A, McDonald TO, Sei E, Shi X, Wang Y, et al. Punctuated copy number evolution and clonal stasis in triple-negative breast cancer. Nat Genet. 2016;48:1119–30.

29. Baslan, T. et al. Genome-wide copy number analysis of single cells. Nat. Protoc. 7, 1024–1041 (2012).

30. McKenna, A. et al. The Genome Analysis Toolkit: A MapReduce framework for analyzing next-generation DNA sequencing data. Genome Res. 20, 1297–1303 (2010).

31. Lodato, M. A. et al. Somatic mutation in single human neurons tracks developmental and transcriptional history. Science (80-.). 350, 94–98 (2015).

32. Koboldt, D. C. et al. VarScan 2: Somatic mutation and copy number alteration discovery in cancer by exome sequencing VarScan 2: Somatic mutation and copy number alteration discovery in cancer by exome sequencing. Genome Res. 22, 568–576 (2012).

33. Cibulskis, K. et al. Sensitive detection of somatic point mutations in impure and heterogeneous cancer samples. Nat. Biotechnol. 31, 213–219 (2013).

