## Supplementary figures and images for "Evaluating Tumor Evolution via Genomic Profiling of Individual Tumor Spheroids in a Malignant Ascites from a Patient with Ovarian Cancer Using a Laser-aided Cell Isolation Technique"

### Supplementary Materials

Additional file1: Figure S1

**Overall process for detecting single nucleotide variants**

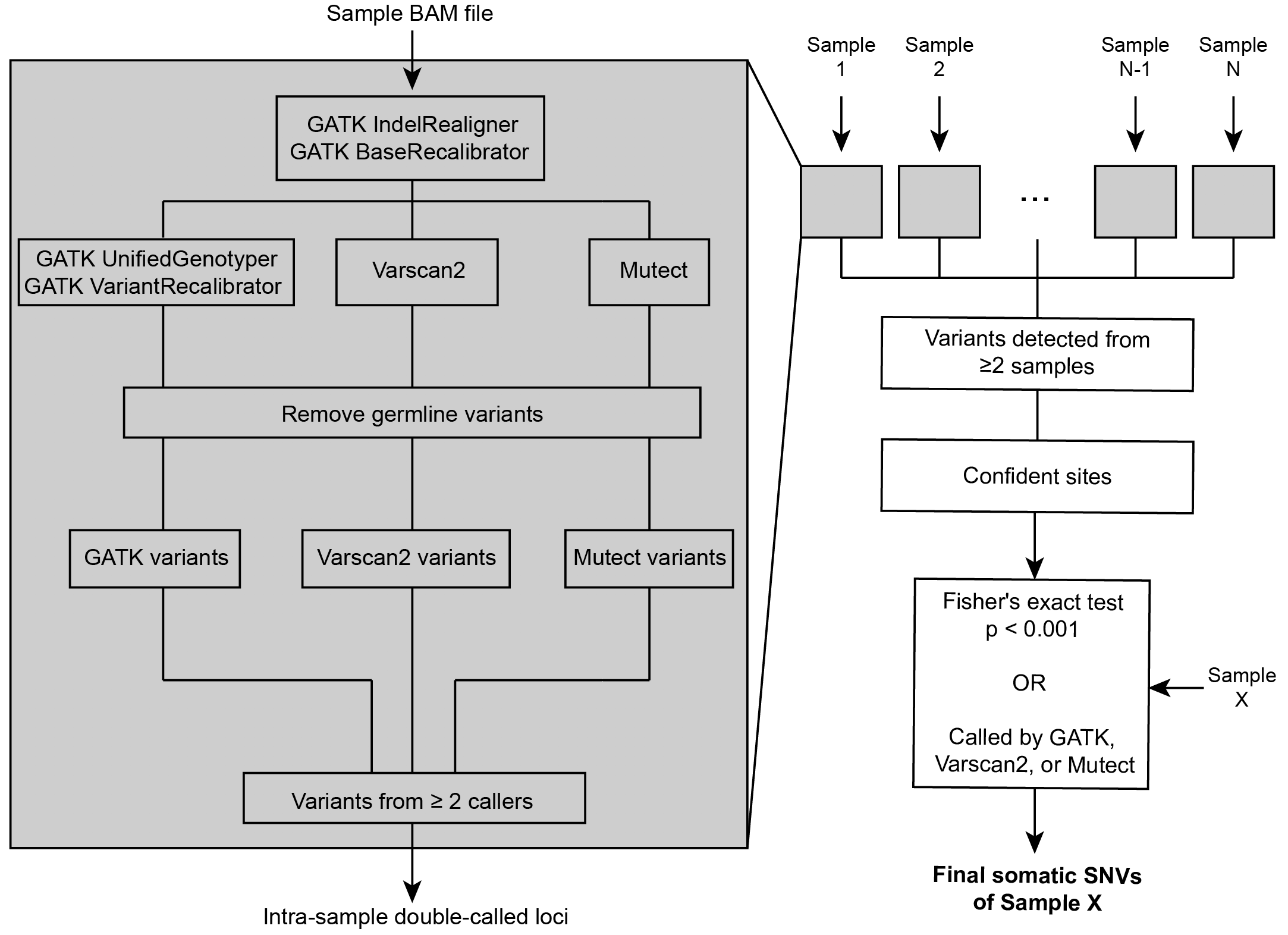
